# A browser-based platform for storage, visualization, and analysis of large-scale 3D images in HPC environments

**DOI:** 10.1101/2024.10.04.616591

**Authors:** Logan A Walker, Wei Jie Lee, Bin Duan, Menelik Weatherspoon, Ye Li, Fred Y. Shen, Mou-Chi Cheng, Xiaoman Niu, Jacob Eggerd, Bin Xie, Jimmy Hon Pong Cheng, Humphrey Xu, Xiao-Xiang Zhong, Hehai Jiang, Meng Cui, Yan Yan, Dawen Cai

## Abstract

High-throughput microscopy necessitates 3D image storage, visualization, and analysis in terabyte to petabyte scales. Centralized high-performance computing (HPC) infrastructure provides the resources, but interfacing software is limited. We developed an extensible platform with a scalable image storage format (SISF) and content delivery network (CDN) to enable fast, random access to compressed data. Additionally, we built a cloud-based nTracer2 software to annotate neuron morphology from SISF images in a web browser.

---

Light-sheet and other high-throughput microscopy techniques have now become broadly accessible to research labs, yet their applications are typically bottlenecked by the data storage and analysis ability of the single imaging computer^1–3^. Institutional HPC infrastructures can handle the increasing amount of imaging data produced within a single experiment beyond the terabyte scale^4,5^. However, software tools that allow efficient interfacing with HPC data storage systems are required for interactive visualization, annotation, and parallel analysis of large multidimensional microscopy datasets.

With careful consideration of the data organization on HPC systems, we have defined a novel Scalable Image Storage Format (SISF; **Fig. 1a**). SISF datasets are hierarchically split into small unit chunks based on voxel resolution and spatial location. Collections of chunks are grouped into “shards” as individual files on the storage system. The two-level indexing SISF system allows retrieving arbitrary subvolumes while alleviating the challenges of storing and accessing many small files, as seen in OME-Zarr format^6^, which can exceed file count quotas on supercomputing systems. Compared to other large-scale image formats (**Supp. Fig. 1**), SISF contains individual shard metadata files defined separately from the global image metadata, allowing an individual shard to be read and parsed independently. Therefore, it can be advantageous to store small metafiles on expensive but fast media, such as SSD and RAM, satisfying frequent access needs, whereas large image files can be stored on HDD or tape to reduce cost. SISF’s unique metadata structure also enables easy manipulation of sub-regions within each shard, which allows a virtual stitching function to hide overlapping regions from a tiled microscopy experiment without trimming those regions from the raw data (**Fig. 1a, inset, Fig 2c**). We implemented an open-source Python and C++ library to access SISF files. We found that SISF’s Python read performance is comparable to Zarr’s when used with SSD storage and exceeds Zarr’s when HDD storage is used (**Fig. 1b, Supp. Fig. 2**). In terms of Python write performance, we found that SISF outperforms Zarr when creating new archives on both SSD and HDD (**Fig. 1c**). To benchmark SISF’s real-world application write speed, we utilized the C++ library for ingesting TIFF images to SISF and recorded a rate of ∼1TB/minute using an SSD storage array on the University of Michigan supercomputer, which is ten times faster than the data production rate of any published microscope. Finally, the optimized HPC I/O performance permits us to implement lossless (Zstd) and lossy (H.264) compressions to save storage space and achieve real-time decompression for visualization, based on prior work^7,8^.

**Fig. 1.**
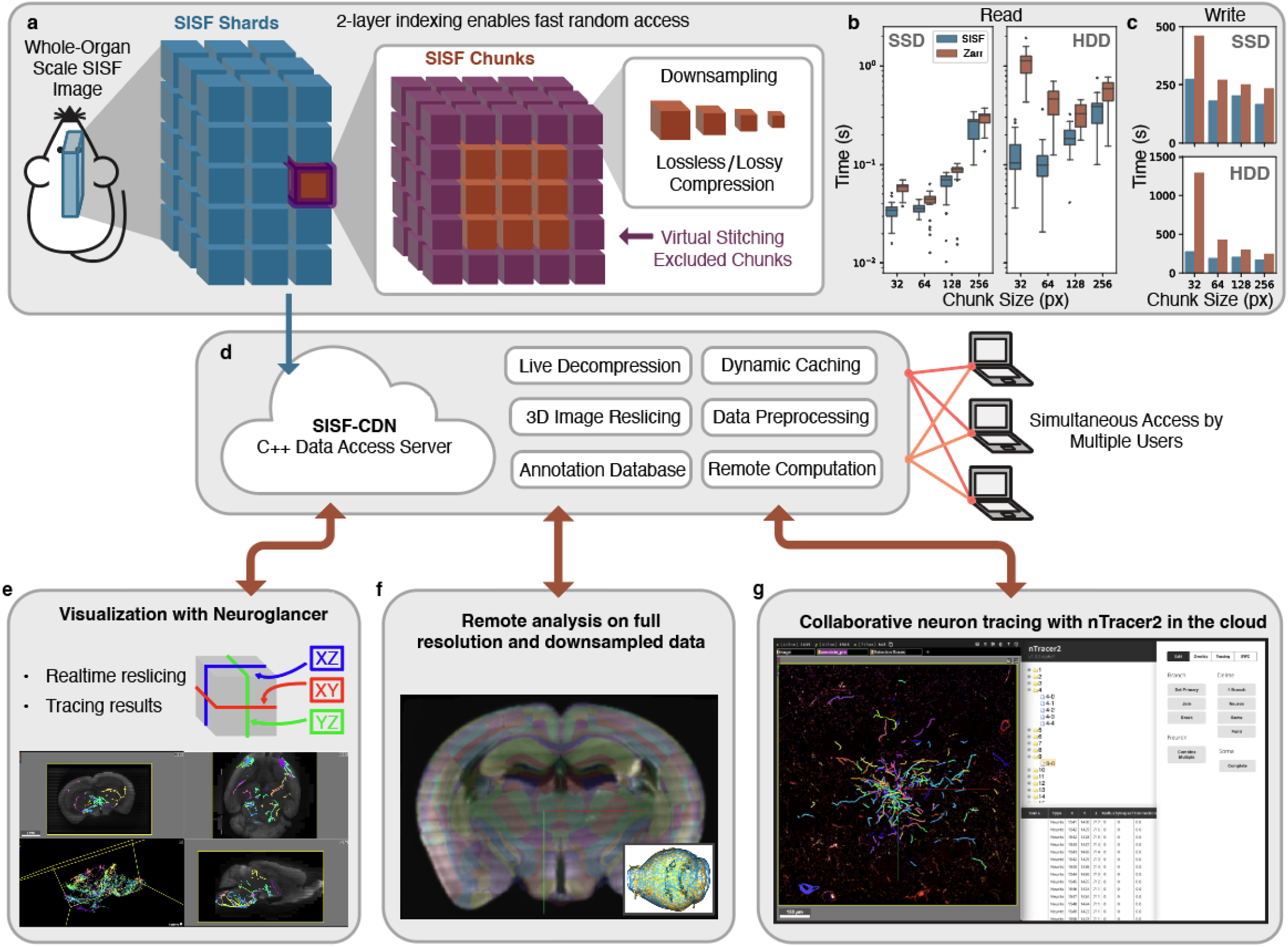
An overview of the SISF-CDN / nTracer2 platform. **a**, A diagram of the SISF file structure. Each dataset is stored in multiple resolutions and split into two layers of chunking to allow quick lookup. Each data chunk can be compressed using ZSTD or H.264 and can be virtually stitched to account for imaging tile overlaps; **b**, The time to read a 256×256×1 image tile from a SISF file compared to Zarr at different storage chunk sizes; **c**, Measurement of the time required to create a 29GB dataset using SISF and Zarr. **d**, an overview of the structure and features of the SISF-CDN. **e-g**, example use cases for the nTracer2 software package; **e**, SISF-CDN converts and reslices 3D images in real time for orthogonal view visualization; **g**, Remote data analysis of images stored by the CDN is enabled by API access; **f**, nTracer2 allows 3D neuron morphology to be reconstructed simultaneously by multiple users from images stored in the CDN. SSD, solid state-based storage; HDD, hard drive-based storage.

**Fig. 2.**
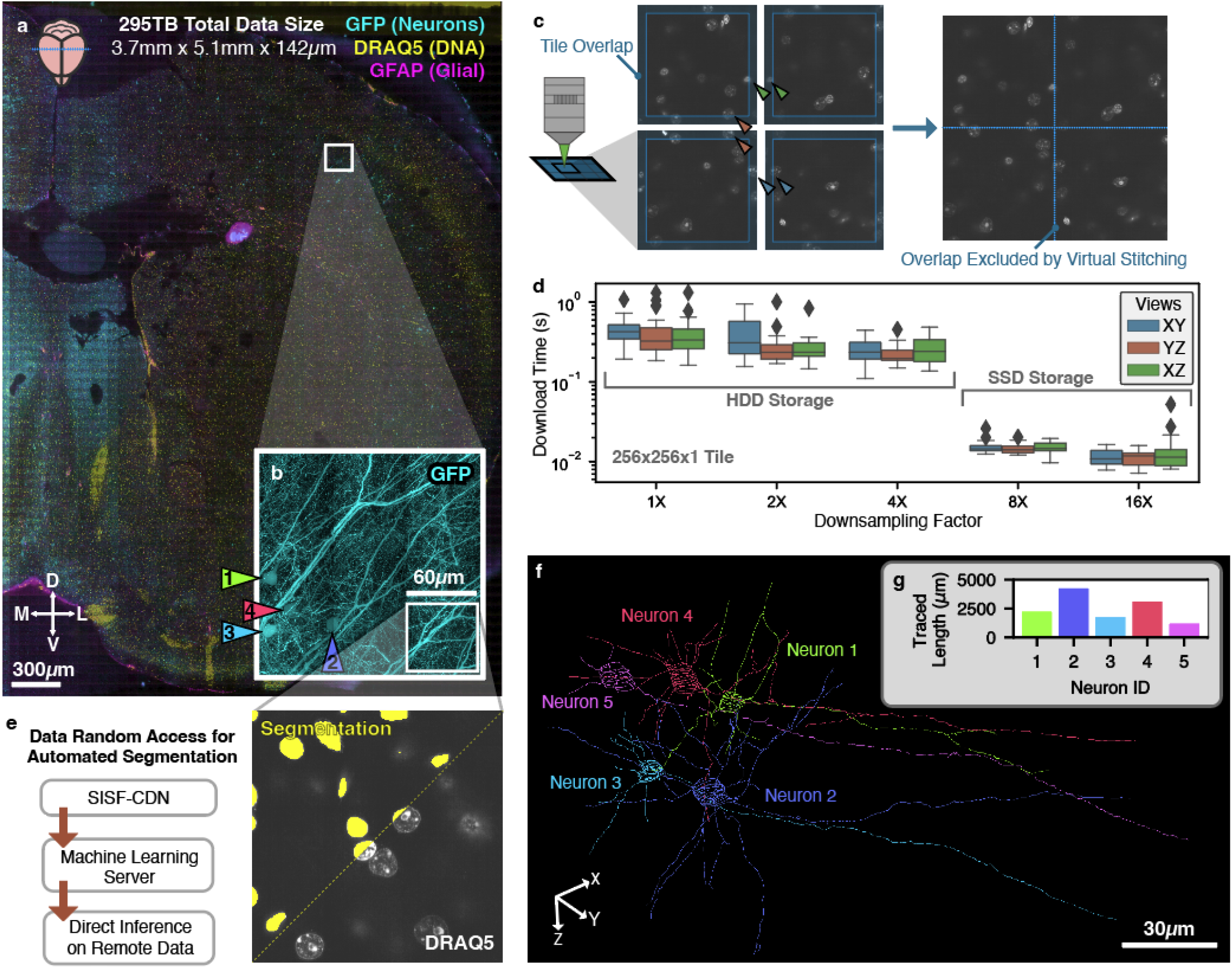
SISF-CDN / nTracer2 enables full-stack analysis of petabyte-scale images. **a**, an overview of a 108,000 × 172,000 × 2,000 voxel (XYZ, 30 × 30 × 70 nm^3^ voxel size) image of a GFPm mouse brain slice labeled with DRAQ5 and GFAP. **b**, magnified view, showing a maximum projection of the GFP-labeled neurons along the transverse plane. Arrowheads indicate four cell bodies; **c**, Schematic showing how the SISF “virtual stitching” feature allows overlapping regions to be excluded from the visualization; **d**, Measurement of download time required to retrieve resliced orthogonal views; **e**, SISF-CDN was used to provide data directly to a machine learning server running an nnU-Net for nuclei segmentation by multiple GPUs in parallel. **f**, Five neurons, including the four identified in **b**, were reconstructed using nTracer2 by multiple users in different locations; **g**, quantification of the tracing length of each neuron in **f**. Scale bars are calibrated for an estimated 3.5X expansion factor.

To facilitate programmatic accessing SISF data files over the local network or Internet, we have created the SISF-CDN server application (**Fig. 1d**). SISF-CDN can manage petabyte-scale datasets and access through a library of Application Programming Interface (API) functions. Compared to direct HTTP access that downloads whole raw data chunks^6^, our CDN approach allows data reslicing (**Fig. 1e**) and downloads only the requested specific subsets at hundreds of megabytes per second. Notably, the computational workload is handled by the server, reducing the hardware requirements for remote clients. As the API is compatible with the naming schema used by the Google Neuroglancer^9^ precomputed image format and related tools^10,11^, images stored on the SISF-CDN can be visualized interactively in a web browser for distributing large-scale images to the public (**Fig. 1e**). The ability to access arbitrary 3D subvolumes remotely in defined resolutions also allows using centrally-stored data to develop analysis pipelines, e.g., registration of downsampled whole mouse brain images to the Common Coordinate Framework^12^ (CCF; **Fig. 1g, Supp. Fig. 3**).

In addition to image data, we have implemented the ability to store and manipulate image annotations by the CDN. For instance, neuron tracing results are stored in a custom SQLite3 schema, based on the SWC file format for reconstructing neuron morphology^13^, to be edited in real-time by multiple users and parsed faster than native text files. This makes the CDN a valuable hub for human and automated analysis from remote computing systems. Building on previous work^14^, we developed a React-based web application, nTracer2, which allows multiple users to simultaneously annotate the same image in Neuroglancer (**Fig. 1f, Supp. Fig. 4**). As the semi-automated tracing computation is directly provided by the CDN, nTracer2 annotation is performed entirely in the cloud without leaving the HPC environment where data is stored.

To demonstrate the use of nTracer2 as a full-stack software solution for neuron reconstruction experiments, we prepared a 0.5 mm slice of brain tissue from a Thy1-GFP line M^15^ mouse with a modified miriEx^16^ expansion microscopy protocol, which was then imaged with a custom line confocal imaging system (**Fig. 2**). The complete image was 295TB of raw data, forming a 3-channel volumetric dataset of 108,000 × 172,000 × 2,000 voxels (37 teravoxels) per channel. **Fig. 2a** and **2b** show the overview and magnified view of two channels containing DRAQ5-labeled nuclear DNA (yellow), GFP-labeled neurons (cyan), and GFAP-labeled glial cells (magenta), respectively. This image was converted into a SISF dataset, where the virtual stitching feature was used to remove the overlapping regions between imaging tiles (**Fig. 2c**). We benchmarked the real-world data access time and found that SISF-CDN can deliver all three orthogonal views to clients at a consistent speed over the network (**Fig. 2d, Supp. Fig. 5**). Random data access allows us to download visually-inspected representative subvolumes to train a nnU-Net^17^ model for soma segmentation from the DRAQ5 image channel (**Fig. 2e**). One can then take advantage of the optimized parallel read function of SISF-CDN to inference the whole image data using HPC resources. Finally, neurons were traced using the nTracer2 interface and analyzed using nGauge^18^ (**Fig. 2f-g**).

Together, these results demonstrate SISF-CDN’s unique virtual stitching function and how the nTracer2 platform can be integrated in every step of the bioinformatic analysis pipeline for a neuron morphology experiment. We also demonstrated that the SISF-CDN architecture is versatile to all computational environments, including individual lab servers and large-scale institutional HPC systems.

## Acknowledgments

The authors thank the Brain Image Library (BIL) staff for assistance in testing nTracer2 at the Pittsburgh Supercomputing Center (PSC). The authors thank Brock Palen (Michigan Advanced Research Computing) for coordination efforts and assistance in testing nTracer2 on the University of Michigan research infrastructure. LAW was supported by the University of Michigan Rackham Predoctoral Fellowship. This work was funded by the National Institutes of Health (NIH) Brain Research Through Advancing Innovative Neurotechnologies (BRAIN) initiative grants RF1MH123402 and RF1MH133764 to DC and YY, RF1MH124611 to DC, MC, and YY, and the National Science Foundation grant NSF-1707316 to DC (Neuronex-MINT).

## Author Contributions

LAW and DC conceptualized nTracer2 and SISF. LAW, WJL, MW, BD, HX, and XXZ implemented different portions of the code and performed unit testing. FYS, MM, XN, and MCC prepared tissue samples which were used in validation experiments. LAW, WJL, MW, YL, and HPJC performed neuron tracing experiments. LAW, HJ, MC, and DC designed the custom microscope for imaging test data. All authors edited and approved the manuscript, which was drafted by LAW, YL, and DC.

## Competing Interests

Elements of the SISF file format design are subject to US patent application #20240144418 by the University of Michigan, for which LAW and DC are authors. The authors declare no other competing interests.

## Methods

### Mouse Sample Data Preparation

For the data used in benchmarking (**Fig 1b,c, Supp. Fig. 5, Supp. Fig. 6**), a ChAT-IRES-Cre (JAX 006410) mouse was stereotaxically injected with Brainbow AAVs^26^ 2/9 in the dorsal striatum. This results in multicolor fluorescent labeling of most ChAT+ neurons. 200μm sections were antibody amplified and prepared with the miriEx^16^ expansion microscopy protocol, followed by imaging with a Zeiss LSM 780 confocal microscope as described previously. This volume was imaged with estimated XYZ resolution of 100 × 100 × 300 nm and was stitched using BigStitcher^27^ Fiji plugin. This yielded an ∼300 × 300 × 300 µm^3^ image volume with a XYZ resolution of 2897 × 2866 × 1765 px^3^ after stitching. Histogram matching was done to normalize intensity between z-slices in image stacks using the nTracer1 Align-Master ImageJ/Fiji plugin^14^. All data was converted from TIFF into SISF files during the benchmarking described below in Python.

For the whole-slice dataset (**Fig. 2**), a Thy1-GFP line M (JAX 007788) mouse was sacrificed and prepared using a modified miriEx^16^ expansion microscopy protocol. The gel was stained with DRAQ5 (ThermoFisher Scientific) to label DNA, as well as antibody labeling for GFAP (Rabbit anti-GFAP, Dako Z033401-2; Anti-Rabbit AF568, Abcam ab175694). Imaging was performed using a custom line confocal microscope using 488nm, 560nm, and 642nm lasers to image native GFP, GFAP-AF568, and DRAQ5, respectively. This yielded 4,644 overlapping image tiles, each of which was 2304 × 2304 × 2000 voxels in size (XYZ). These were combined into one SISF image of 108,000 × 172,000 × 2,000 voxel with a 105 × 105 × 250 nm^3^ imaging voxel size. We estimated that the sample was expanded by 3.5X, which means the image had an effective ∼30 × ∼30 × ∼70 nm^3^ voxel size. All scale bars for this data were scaled by this voxel size so that the figures represent the size of the original tissue. The virtual stitching feature of SISF-CDN was used to exclude a 304px overlap in the x and y dimensions, forming a continuous image. Notably, virtual stitching enables the retention of overlap in an accessible manner for downstream analysis.

### SISF File Organization

As shown in **Supp. Fig. 1a-c**, SISF files contain three parts: a metadata file (“metadata.bin”) which defines the size of the global image, a folder of shard data (“data”), and a folder of metadata (“meta”). The global metadata file stores a packed description of the attributes of the entire image, including the voxel size, channel count, and optical resolution. The files stored in the “data” and “meta” folders each represent a shard that is named according to the coordinates in 3D space, the channel number, and the downsampling factor (1X, 2X, etc.). Because each shard is large (typically a 1000-3000px cube; however, this is configurable), the data is further split into small chunks and sharded into one continuous data file. The meta file for each shard provides a lookup table for where each chunk in the data file is located and its respective size (which can vary due to compression). The data inside the chunks can be compressed using a variety of methods such as Zstd^23^ or H.264^24^ (video compression mode), motivated by previous work^7^. This hybrid structure allows SISF to be flexible in a variety of deployments, which can be optimized in different ways.

For each shard, SISF allows a crop factor to be defined. This allows shards to contain a larger amount of data than the region which it represents. The canonical use of this feature is the implementation of translational tile alignment that can be performed inside SISF without deleting the overlapping image region, which we have termed “virtual stitching”.

We have created a Python library pySISF [https://github.com/Cai-Lab-at-University-of-Michigan/pySISF] for the creation and reading of SISF files, which is available on the Python pip repository. pySISF natively supports the parallel creation of SISF files and uses the Numba^25^ compiler to accelerate tasks such as image downsampling. A C++ library is also available to accelerate the reading of SISF files. We have created example scripts for converting input file types, such as TIFF series, into SISF files. Additionally, SISF can be easily integrated into custom microscopy pipelines where individual tile scans are saved as SISF shards.

### SISF vs. Zarr Benchmark

All benchmarking was performed on a custom Linux server with two EPYC 7551 CPUs, 512GB of DDR4 RAM, and a 40 Gbps fiber optic network connection running Ubuntu 20.04LTS. During all tests, only the benchmarking task was run to reduce potential artifacts from other users. SSD tests were performed using a ZFS RAID-Z1 array of five PNY CS2241 4TB NVMe SSDs mounted locally to the test system. HDD tests were performed using hard drive-based HPC storage hosted by the University of Michigan Advanced Research Computing (ARC) mounted over Network File System (NFS) v4 to the host system. This allows us to simulate the latency of a typical high-performance computing storage environment.

To compare the performance of SISF and Zarr (**Fig 1b,c, Supp. Fig. 2**), one channel of the ChAT Brainbow image described above was used for a total raw data size of 29.31GB. Before performing all benchmarking tests, the Linux server disk cache was completely cleared to prevent caching from impacting benchmark times and reduce bias introduced by caching of volume metadata. The raw image was then converted into SISF and Zarr archives of isotropic chunk sizes of 32, 64, 128, and 256 compressed with Zstd^23^ level 5, and the write times were recorded for each setting. From these files, N=30 random coordinates were generated for each chunk size, and a ZXY chunk of 1 × 256 × 256 was read and timed at each coordinate using a Python script. This test was permuted for ZXY chunks of 256 × 1 × 256 and 256 × 256 × 1 to ensure no bias for dimension selection. Reads were selected so as not to fall on file chunk boundaries and to avoid increasing the average number of chunks that needed to be read. Python v3.11 and Zarr 2.14.2 were used for this analysis.

### SISF-CDN Implementation

The SISF-CDN is implemented in C++ and compiled using GNU C++ compiler 9.4.0. The Crow C++ library V1.0+5 (https://crowcpp.org/) was used to implement the CDN HTTP functionality, and the SQLite v3 (https://www.sqlite.org/) library is used to store SWC tracing data on the CDN. Neuron SWC manipulation is performed with a novel library based on nGauge^18^, which has been translated to C++ to allow fast processing of hundreds of neurons at a time. We modeled the CDN API to match the neuroglancer “precomputed” format structure for easy cross-compatibility without requiring any client code changes. API endpoints are implemented for SWC editing, uploading, and access to enable. A Python API was implemented to allow direct data download from the CDN into different Python scripts, and it is used in the nTracer2 backend.

### nTracer2 Tracing Application Implementation

The nTracer2 application server and UI are implemented using a collection of modern web development frameworks. This includes React [https://react.dev/] and jsTree [https://www.jstree.com/] for the UI and Flask [https://flask.palletsprojects.com/] for the server backend. The data viewer uses Neuroglancer^9^, with additional UI elements and shaders added via Python scripts. All builds are managed with Poetry [https://python-poetry.org/] to provide reproducible Python environments. Parts of the Python server backend are implemented in Cython [https://cython.org/], which allows Python-compatible C code to be used to accelerate the image analysis. Neuron SWC manipulation is performed using nGauge^18^. The nTracer2 application and UI are provided as a Docker container for easy deployment and management in end-user systems, as detailed below.

### CDN SWC Serialization

In typical applications, SWC files are stored as text files for ease of portability^13^. When developing nTracer2, we found this is limiting because of the need to rapidly decode and edit SWC files during tracing. As such, we implemented a C++ library that allows the storage of SWC files inside SQLite3 database files. Briefly, this conversion is done as follows: inside the database file for each image, two tables are created. One table contains the metadata for each neuron, including each neuron’s identifier ID, the coordinates of the neuron’s soma, and metadata about the inferred cell type. The second table stores the data traditionally found in a SWC file, namely a linked list of all points making up the neuron. This structure allows neurons to be quickly identified using SQL queries of spatial coordinates, as well as visualized using a live conversion performed by the CDN to generate a binary trace representation.

### Docker Usage

Docker^28^ is a widely used development platform for creating reproducible and reliable containerized infrastructure projects. To facilitate the usage and deployment of nTracer2, we have produced two docker images: one that hosts the SISF-CDN and one that hosts the nTracer application server. These docker images allow the nTracer2 applications to be deployed quickly and easily on different server environments. The CDN docker image can be found on DockerHub at [https://hub.docker.com/r/geeklogan/sisf_cdn], and the nTracer2 docker image can be found on DockerHub at [https://hub.docker.com/r/geeklogan/ntracer2]. A tutorial on Docker usage can be found on the respective GitHub pages (See **Data and Code Availability**).

### Public Data Visualization

To validate the performance of the SISF-CDN, we accessed a previously-published^29^ green fluorescent protein (GFP) fluorescence micro-optical sectioning tomography (fMOST)^2^ brain dataset, which consists of a ZXY size of 11,464 × 20,821 × 30,801 voxels (14.7TB of raw data). This data represents a Fezf2-CreER/wt;Ai166(TIT2L-MORF-ICL-tTA2)/wt animal and was accessed from the Brain Image Library (BIL; https://www.brainimagelibrary.org/; https://doi.org/10.35077/ace-add-lit). Additionally, 13 large SWC neuron reconstructions were uploaded to the CDN. This dataset is shown, as accessed through the SISF-CDN, in **Fig. 2f** and **Supp. Fig. 4**.

### Brain Image to CCF Registration Method

To register a brain image to the mouse Common Coordinate Framework^12^, we implemented a novel warping technique (**Fig. S1a**). First, the input image is resampled to match the resolution of the CCF map (10μm). Next, an artifact removal step removes imaging artifacts, such as scan lines from the resampled input image. The contour of each imaging slice is segmented and used to generate a point cloud representing the outer surface of the brain (**Fig. S1b**). The cortex regions of both images are aligned together first, followed by a global linear alignment. Next, a local nonlinear deformation is used to match parts of the brain that are different shapes relative to the CCF reference. Finally, a deformed CCF image is generated based on this final deformation, which overlays the original data well. This allows the CCF annotation to be overlaid on visualized datasets for identifying specific brain regions and coordinates in different image modalities. All code is implemented in Python. Registering a whole-brain fMOST dataset took ∼50 minutes on a 2.1GHz AMD Ryzen 1950X CPU with 72 GB of memory peak usage. The result of aligning the CCF to the fMOST dataset described above is displayed in **Fig. 2c**, with a larger example provided in **Fig. S1c**.

### CDN Access Benchmarks

To test the access speed of our CDN system when hosted on an HPC infrastructure, we deployed the nTracer2 CDN onto a virtual machine hosted in the University of Michigan ITS data center. Hard drive-based HPC storage hosted by the University of Michigan Advanced Research Computing (ARC) was used to store the test datasets and mounted over NFS v4 over 100Gb ethernet networking. 16× and 32× tiles were stored on a separate caching SSD for accelerating overview tiles. From a desktop machine located in a lab at the University of Michigan, benchmarking was performed by randomly generating N=100 coordinates for each downsampled resolution of the example fMOST image^29^ (XYZ size 31744 × 21504 × 13312) and reading an XYZ 256 × 256 × 1 image tile from the CDN over HTTP using our custom Python library for interacting with the CDN. This analysis was performed with Python 3.10 and the Python requests 2.31.0 library.

### SISF Conversion Benchmark

Data was stored in four IME240 data storage servers (DDN Storage, Los Angeles, CA) located at the University of Michigan Advanced Research Computing (ARC) Great Lakes supercomputer. When combined, these servers provide 80TB of available SSD storage using non-volatile memory express (NVMe) and are connected through 800 Gbps InfiniBand networking. ∼480 Intel Xeon cores were used simultaneously across 10 supercomputer nodes, and timing was performed by parsing workload manager (SLURM) log outputs.

### Data and code availability

All data produced in this study is available from the corresponding author upon request. All code and documentation for nTracer2 is available at https://github.com/Cai-Lab-at-University-of-Michigan/nTracer2 ^30^. The code and documentation for the SISF-CDN is available at https://github.com/Cai-Lab-at-University-of-Michigan/SISF_CDN. The SISF Python library is released at https://github.com/Cai-Lab-at-University-of-Michigan/pySISF. Brain images-to-CCF registration codes are available at https://github.com/Cai-Lab-at-University-of-Michigan/CCF_Registration_Pipeline. Microscopy data is being uploaded to the BIL for public access. The nTracer2 and SISF-CDN codes are available as a Docker image on DockerHub.

## References

1. Voleti, V. et al. Real-time volumetric microscopy of in vivo dynamics and large-scale samples with SCAPE 2.0. Nat. Methods 16, 1054–1062 (2019).

2. Wang, X. et al. Chemical sectioning fluorescence tomography: high-throughput, high-contrast, multicolor, whole-brain imaging at subcellular resolution. Cell Rep. 34, 108709 (2021).

3. Glaser, A. K. et al. A hybrid open-top light-sheet microscope for versatile multi-scale imaging of cleared tissues. Nat. Methods 19, 613–619 (2022).

4. Kenney, M. et al. The Brain Image Library: A Community-Contributed Microscopy Resource for Neuroscientists. bioRxiv 2023.12.22.573024 (2024) doi:10.1101/2023.12.22.573024.

5. Towns, J. et al. XSEDE: Accelerating scientific discovery. Comput. Sci. Eng. 16, 62–74 (2014).

6. Moore, J. et al. OME-NGFF: a next-generation file format for expanding bioimaging data-access strategies. Nat. Methods 18, 1496–1498 (2021).

7. Walker, L. A., McGlothlin, M., Li, Y. & Cai, D. A Comparison of Lossless Compression Methods in Microscopy Data Storage Applications. bioRxiv 2023.01.24.525380 (2023).

8. Duan, B. et al. Artifact-minimized high-ratio image compression with preserved analysis fidelity. bioRxiv 2024.07. 17.603794 (2024) doi:10.1101/2024.07.17.603794.

9. Creators Jeremy Maitin-Shepard Alex Baden1 William Silversmith2 Eric Perlman3 Forrest Collman4 Tim Blakely5 Jan Funke Chris Jordan Ben Falk6 Nico Kemnitz7 tingzhao Chris Roat Manuel Castro Sridhar Jagannathan8 moenigin Jody Clements Austin Hoag9 Bill Katz10 Dave Parsons Jingpeng Wu11 Lee Kamentsky Pavel Chervakov Philip Hubbard12 Stuart Berg John Hoffer13 Akhilesh Halageri Christian Machacek Kevin Mader Lutz Roeder14 Peter H. Li Show affiliations 1. @omnisci 2. Seung Lab @ Princeton University 3. Yikes LLC 4. Allen Institute for Brain Science 5. @google 6. @stsci 7. @seung-lab @zettaai 8. Charité Universitätsmedizin Berlin 9. Princeton University 10. Janelia Research Center 11. @flatironinstitute 12. Howard Hughes Medical Institute, Janelia Research Campus 13. Harvard 14. @microsoft. Google/neuroglancer: (2021). doi:10.5281/zenodo.5573294.

10. Silversmith, W. et al. Igneous: Distributed dense 3D segmentation meshing, neuron skeletonization, and hierarchical downsampling. Front. Neural Circuits 16, 977700 (2022).

11. Cloud-Volume: Read and Write Neuroglancer Datasets Programmatically. (Github).

12. Wang, Q. et al. The Allen Mouse Brain Common Coordinate Framework: A 3D Reference Atlas. Cell 181, 936–953.e20 (2020).

13. Nanda, S. et al. Design and implementation of multi-signal and time-varying neural reconstructions. Sci Data 5, 170207 (2018).

14. Roossien, D. H. et al. Multispectral tracing in densely labeled mouse brain with nTracer. Bioinformatics 35, 3544–3546 (2019).

15. Feng, G. et al. Imaging neuronal subsets in transgenic mice expressing multiple spectral variants of GFP. Neuron 28, 41–51 (2000).

16. Shen, F. Y. et al. Light microscopy based approach for mapping connectivity with molecular specificity. Nat. Commun. 11, 4632 (2020).

17. Isensee, F., Jaeger, P. F., Kohl, S. A. A., Petersen, J. & Maier-Hein, K. H. nnU-Net: a self-configuring method for deep learning-based biomedical image segmentation. Nat. Methods 18, 203–211 (2021).

18. Walker, L. A. et al. nGauge: Integrated and Extensible Neuron Morphology Analysis in Python. Neuroinformatics 20, 755–764 (2022).

19. Moore, J. et al. OME-Zarr: a cloud-optimized bioimaging file format with international community support. Histochem. Cell Biol. (2023) doi:10.1007/s00418-023-02209-1.

20. Bria, A., Iannello, G., Onofri, L. & Peng, H. TeraFly: real-time three-dimensional visualization and annotation of terabytes of multidimensional volumetric images. Nat. Methods 13, 192–194 (2016).

21. Sharding codec (version 1.0) — Zarr specs documentation. https://zarr-specs.readthedocs.io/en/latest/v3/codecs/sharding-indexed/v1.0.html.

22. Kent, B. R. 3D Scientific Visualization with Blender. (Morgan & Claypool Publishers, 2014).

23. Collet, Y. & Kucherawy, M.S. RFC 8878: Zstandard Compression and the ‘application/zstd’ Media Type. https://www.rfc-editor.org/rfc/rfc8878.

24. Bross, B., Chen, J., Ohm, J.-R., Sullivan, G. J. & Wang, Y.-K. Developments in International Video Coding Standardization After AVC, With an Overview of Versatile Video Coding (VVC). Proc. IEEE 109, 1463–1493 (2021).

25. Lam, S. K., Pitrou, A. & Seibert, S. Numba: a LLVM-based Python JIT compiler. in Proceedings of the Second Workshop on the LLVM Compiler Infrastructure in HPC 1–6 (Association for Computing Machinery, New York, NY, USA, 2015).

26. Cai, D., Cohen, K. B., Luo, T., Lichtman, J. W. & Sanes, J. R. Improved tools for the Brainbow toolbox. Nat. Methods 10, 540–547 (2013).

27. Hörl, D. et al. BigStitcher: reconstructing high-resolution image datasets of cleared and expanded samples. Nat. Methods 16, 870–874 (2019).

28. Boettiger, C. An introduction to Docker for reproducible research. Oper. Syst. Rev. 49, 71–79 (2015).

29. Peng, H. et al. Morphological diversity of single neurons in molecularly defined cell types. Nature 598, 174–181 (2021).

30. Walker, L. & Lee, W. J. Cai-Lab-at-University-of-Michigan/nTracer2: First Release. (Zenodo, 2024). doi:10.5281/ZENODO.13697099.

