## Supplemental Figures 1-5 for "A browser-based platform for storage, visualization, and analysis of large-scale 3D images in HPC environments"

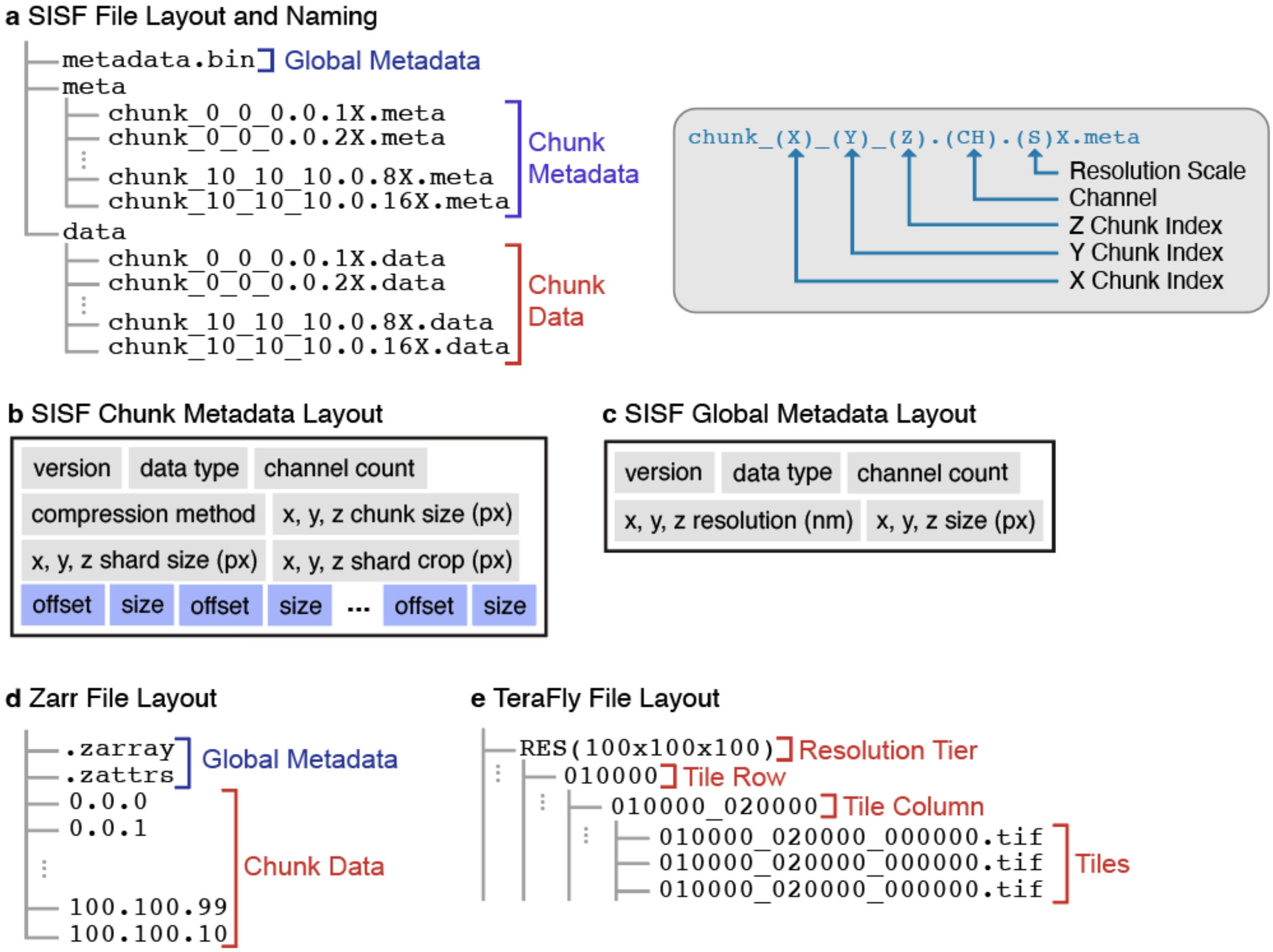


**Supp. Fig. 1 | The SISF file format and comparison to other large-scale image formats. a,** The layout of the data in a SISF file. Folder structure is indicated using indentation and drawn using a tree structure. See Online Methods for a detailed description of the layout; **b**,**c** Diagrams of the layout of the binary global and chunk metadata. Each block is one header element stored using a packed binary format; **d**,**e** example layouts of Zarr[19] and TeraFly[20] files, respectively, when using the default settings. Notably, SISF uses shards not present in Zarr v2 and TeraFly files. At the time of submitting this manuscript, a proposed Zarr v3[21] sharded mechanism is in its early beta stage. However, it does not provide the required functionality for a full-stack CDN performance. In addition, its sharding mechanism does not allow critical features such as the virtual stitching provided by SISF.

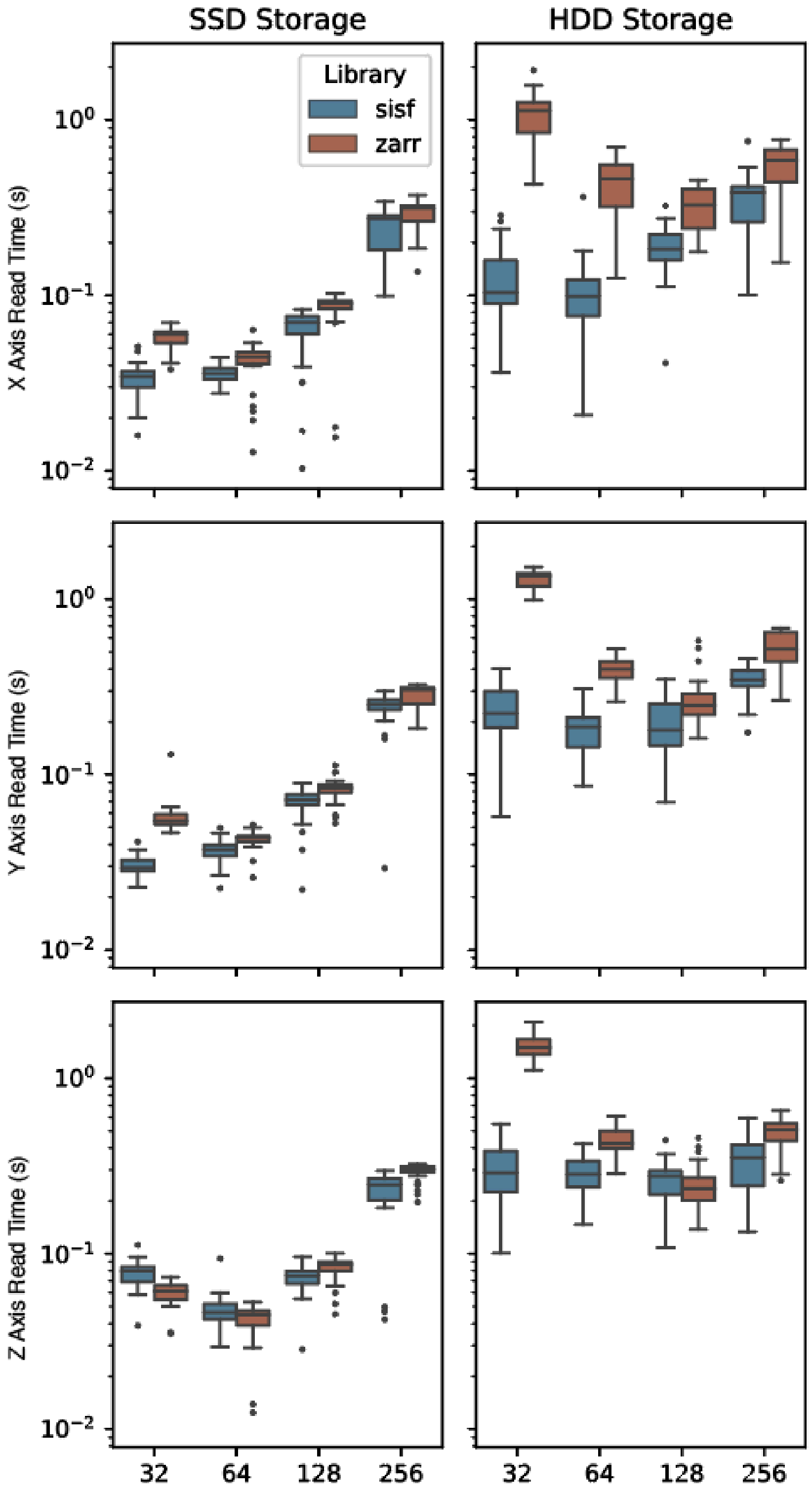


**Supp. Fig. 2 | Benchmarking of SISF and Zarr image access.** An extension of the analysis presented in Figure 1b. Here, each view is permuted, showing that there is little access time bias when accessing frames aligned with different views. In particular, this is 1x256x256 voxel, 256x1x256 voxel, and 256x256x1 voxel.

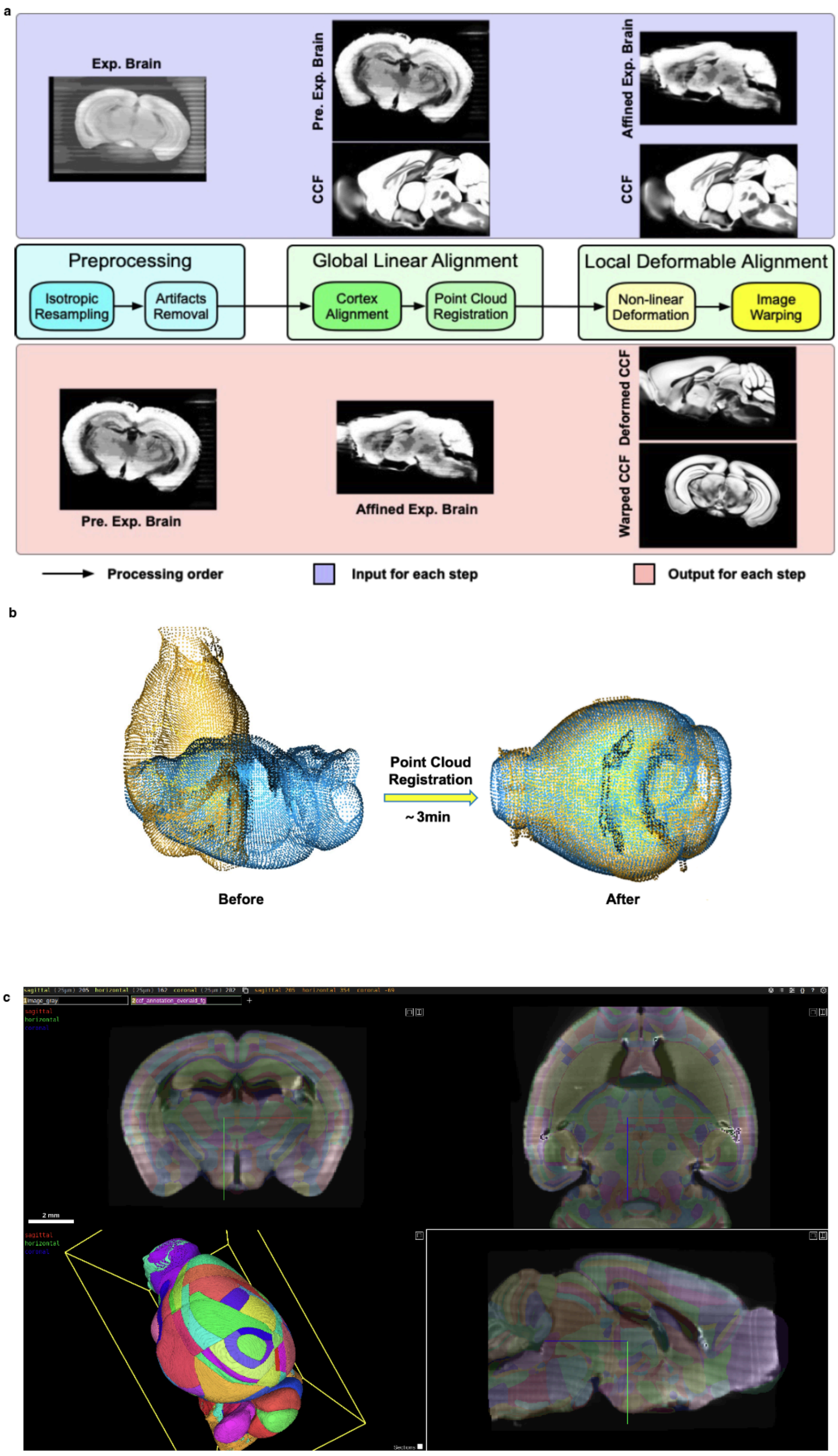


**Supp. Fig. 3 | Registration of the CCF reference atlas to a whole-brain dataset using SISF-CDN.** As an example of the use of SISF-CDN for remote data analysis, we tested a protocol for transforming the Allen Institute Common Coordinate Framework V3[12] (CCF) to match a given image (see Methods). **a**, The workflow implemented for CCF alignment; **b**, An example of the point clouds calculated from the CCF (yellow) and image data (blue); **c**, Plots of the CCF labels in multiple colors overlaid on the image, showing accurate registration.

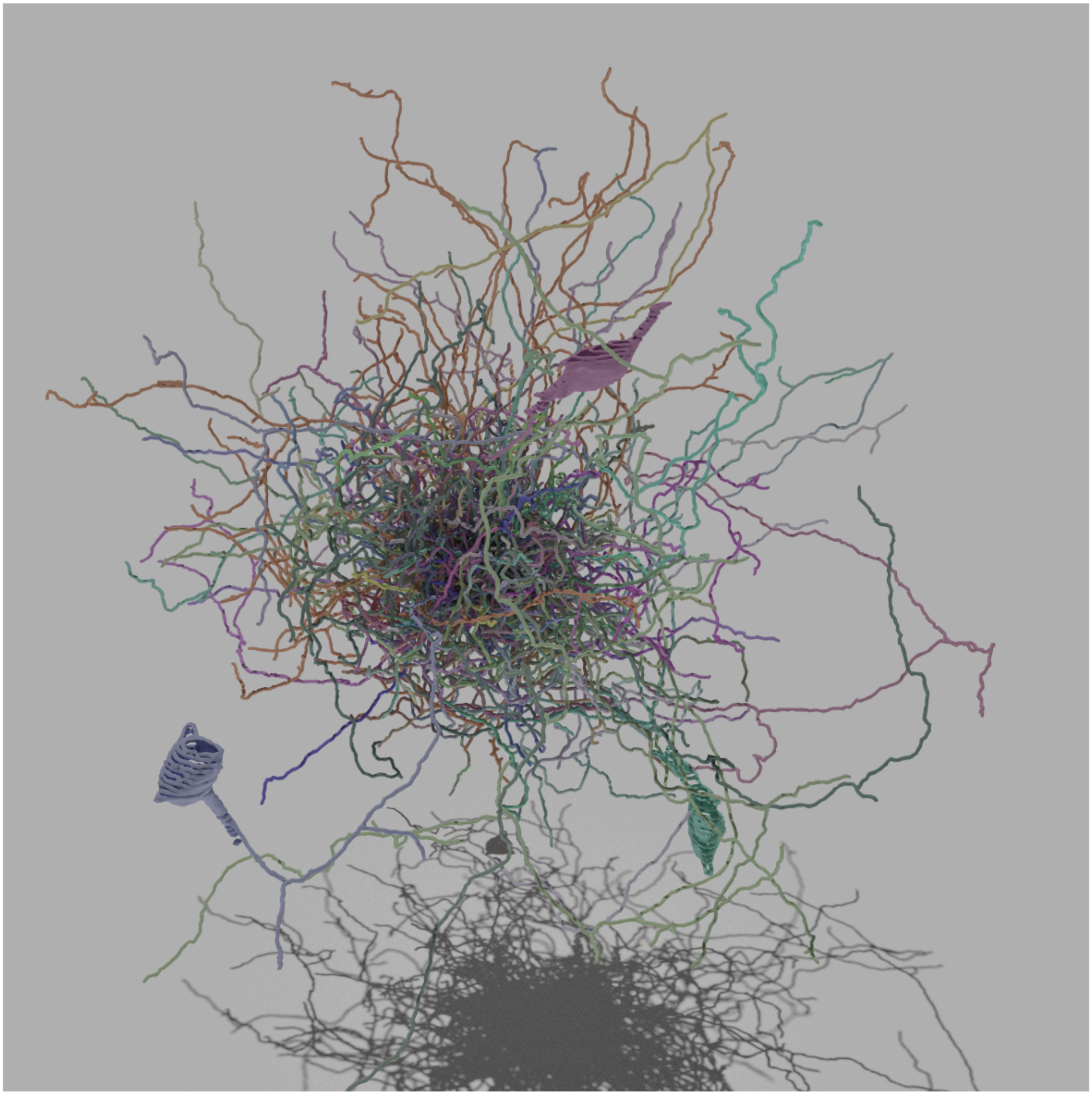

**Supp. Fig. 4 | nTracer2 is capable of multispectral neuronal tracing in a Brainbow labeled mouse brain sample.** We performed a dense reconstruction of a small volume of Brainbow image data, where all the neurons present in the center 512 x 512 x 512 px of the image were fully tracked to the periphery of the image. The 358 reconstructed neurites were rendered using nGauge[18] and Blender[22].

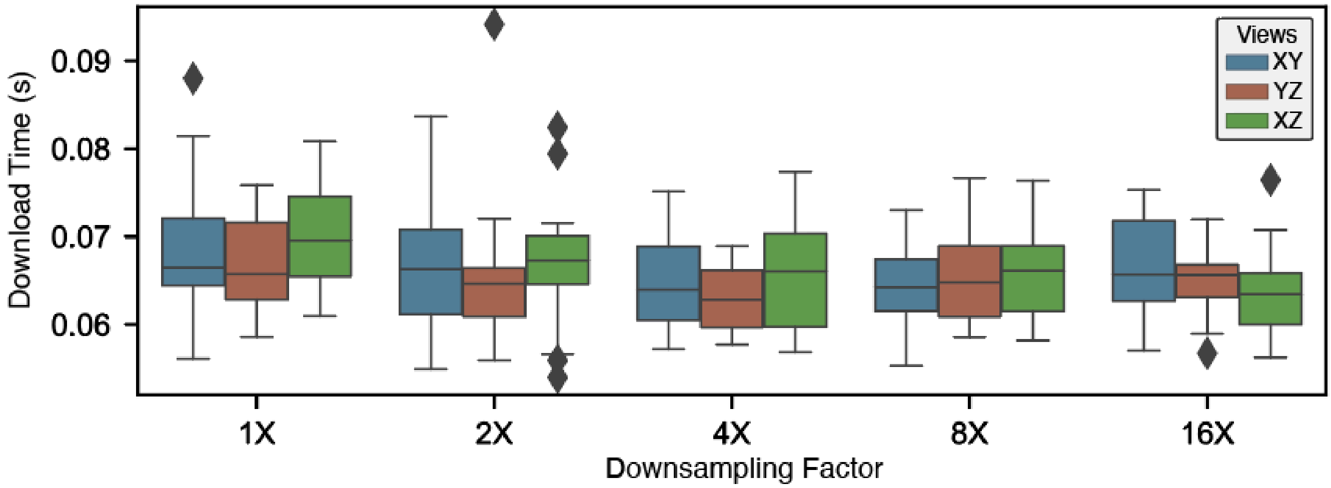


**Supp. Fig. 5 | Additional benchmarking of SISF-CDN image access performance.** The analysis in **Fig. 2d** was repeated for a ~8TB dataset stored exclusively using SSD storage.
